# Potential Opportunity of Antisense Therapy of COVID-19 on an in Vitro Model

**DOI:** 10.1101/2020.11.02.363598

**Authors:** A.N. Goryachev, S.A. Kalantarov, A.G. Severova, A.S. Goryacheva

## Abstract

Data on potential effectiveness and prospects of treatment of new coronavirus infection of COVID-19 caused by virus SARS-CoV-2 with the help of antisense oligonucleotides acting against RNA of virus on an in vitro model are given. The ability of antisense oligonucleotides to suppress viral replication in diseases caused by coronaviruses using the example of SARS and MERS is shown. The identity of the initial regulatory section of RNA of various coronaviruses was found within 50 - 100 nucleotides from the 5’-end, which allows using antisense suppression of this RNA fragment. A new RNA fragment of the virus present in all samples of coronovirus SARS-CoV-2 has been identified, the suppression of which with the help of an antisense oligonucleotide can be effective in the treatment of COVID-19. The study of the synthesized antisense oligonucleotide 5’ - AGCCGAGTGACAGCC ACACAG, complementary to the selected virus RNA sequence, was carried out. The low toxicity of the preparations of this group in the cell culture study and the ability to reduce viral load at high doses according to real time-PCR data are shown. The cytopathogenic dose exceeds 2 mg / ml. At a dosage of 1 mg / ml, viral replication is reduced by 5-13 times. Conclusions are made about the prospects of this direction and the feasibility of using the inhalation way of drug administration into the body.

## Introduction

The coronavirus pandemic has shown the unpreparedness of the world community for the sixth technological paradigm, which is based on nano- and biomedical technologies. Overcoming the interspecies barrier and the accidental penetration of the wild strain of the coronavirus infection SARS-CoV-2 of bats into the human population caused a pandemic and a severe economic crisis due to anti-epidemic measures.

The need for a quickly response to such large-scale threats in the event of a massive influx of infected people and the threat of even greater contamination of the population forces us to look for ways to urgently treat the flow of patients and virus carriers. Moreover, this applies not only to the current, but also to future epidemics that may be caused by other respiratory viruses - influenza, respiratory syncytial virus, rhinoviruses, paramyxoviruses, etc. One of the possible directions in the therapy of viral infections caused by RNA viruses is therapy with antisense oligonucleotides. These drugs act on unique sites of the virus genome and are a promising trend in genetic technologies.

Presently, there are no specific antivirals for a huge array of viruses. However, methods are being developed for specifically selectively “turning off” RNA, including viral, using antisense oligonucleotides. The principle of action of antisense oligonucleotides consists in administering to the body of the patient a drug containing single-strand DNA chains which is complementary to any site of single-strand DNA or RNA, for example, virus RNA. Complementary binding of DNA of the preparation and RNA of the virus leads to impossibility of transcription and translation of viral RNA and cutting of blocked section of RNA of the virus by RNase H. This stops synthesis of new viral particles and prevents intracellular propagation of the virus [12].

Several antiviral antisense drugs have been marketed, for example Fomivirsen^®^ (Vitraven^®^) - a drug against cytomegalovirus (Novartis), Miravirsen^®^ - a drug for the treatment of hepatitis C (Santaris Pharma). Currently, more than 100 drugs for various diseases based on the use of antisense oligonucleotides are undergoing clinical trials in various phases.

Antisense therapy is a developing strategy for the specific treatment of new and socially significant diseases. The principle of RNA inhibition has previously been studied in vitro to inhibit replication of highly pathogenic RNA-viruses [6, 8].

Thus, given the previous experience of antisense oligonucleotide therapy, it can be assumed that this strategy can be applied as an antiviral drug by binding and cleavage of RNA SARS-CoV-2.

Considering that SARS-CoV-2 is an RNA virus that does not integrate into the host genome, the strategy of using antisense oligonucleotides, according to some authors, can give effective results [1, 9].

Prerequisites for the use of antisense therapy in the treatment of COVID 19 exist. Coronavirus infections have previously caused outbreaks of epidemics. In particular, the outbreak of SARS (severe acute respiratory syndrome) in China in 2002-2003 was caused by coronavirus SARS-CoV, the outbreak of Middle Eastern respiratory syndrome (MERS) was also caused by coronavirus [3, 14].

After the SARS epidemic, a number of authors conducted studies on the effect of antisense therapy on the suppression of coronavirus growth in the tissues under investigation [10, 11, 13]. In work (10), the authors found among several sequences the most effective viral transcription inhibitors, suppressing the reproduction of coronavirus in cells and preventing infection of other cells. These are antisense blockers complementary to the initial nucleotides of viral RNA - the so-called regulatory transcription sequences of TRS (in the terminology of the authors) of the coronavirus strain SARS-CoV-Tor2 (GenBank AY274119). The nucleotide sequence of the antisense preparations examined is shown in Table 1:

**Table 1.**
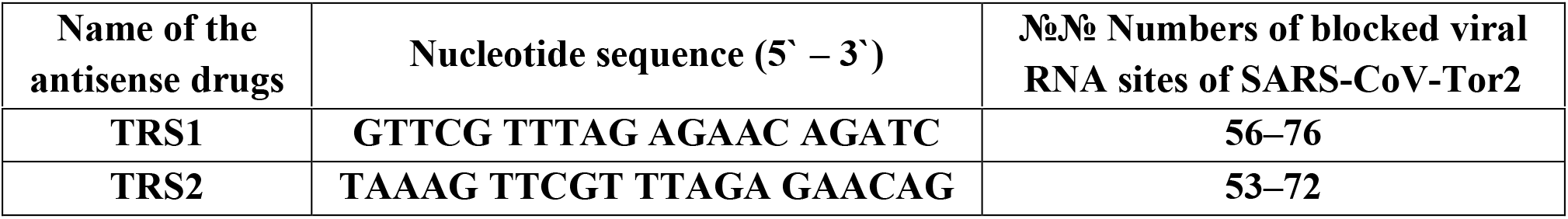
Antisense drugs investigated in work (10)

These sequences are complementary to the following viral nucleic acid sequences (Table 2):

**Table 2.**
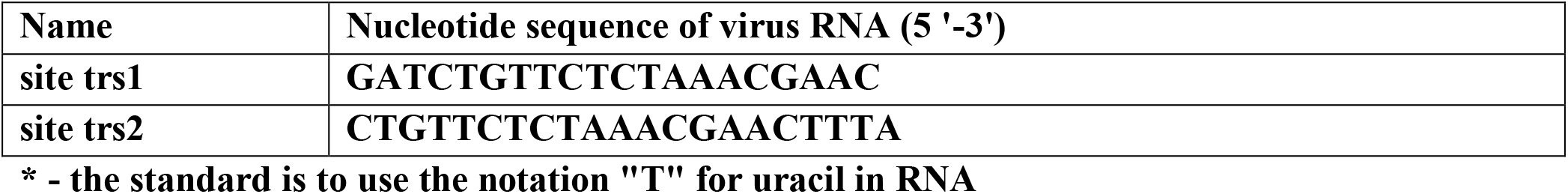
Genetic target sequences of the coronavirus genome for TRS1 and TRS2 drugs.

The authors used morpholine side chain modification to prevent destruction of antisense oligonucleotide and conjugation with arginine polypeptide to improve penetration into infected cells.

By comparison of these nucleotide sequences (table 2) to the sequence of a genome of the SARS-CoV-2 virus (COVID-19) the existence of these sequences almost in all strains of a virus allocated at patients during a pandemic of 2019-2020 was revealed (Appendix 1, the nucleotide sequences are highlighted in red color and a frame).

BLAST analysis (https://blast.ncbi.nlm.nih.gov/Blast.cgi) showed 100% coincidence of the RNA sequence in all coronavirus subtypes, which suggests a high conservatism of this region of the coronavirus genome. This makes it possible to use for the treatment of coronavirus infection those antisense oligonucleotide sequences TRS1 and TRS2, which were studied in 2005 by the collective [10].

However, starting from mid-March 2020, changes have been observed in the virus strains associated with the loss of sites from the 5’ end. In particular, the MT263462 USA: WA 2020-03-23 virus sample did not have the regions designated as trs1 and trs2.

In this regard, the study of an antisense oligonucleotide complementary to the site of the conserved oligonucleotide sequence of the genome of the SARS-CoV-2 virus present in all strains, which was the **goal** of the present study, seems relevant.

In accordance with the purpose of the study, the following tasks were set:

- to select the nucleotide sequence of the virus that is supposed to be inhibited,
- to carry out the synthesis of oligonucleotide,
- to determine cytotoxicity and antiviral activity in an in vitro experiment on cell culture.

## Material and methods

### 1. Selection of nucleotide sequence

The nucleotide sequence that was intended to block in the SARS-CoV-2 virus was selected next to the trs1 and trs2 genes at a distance of four nucleotides from the final region (3’-end) of the trs2 gene. This choice was due to the fact that this sequence enters the site regulating transcription (transcription regulatory site, trs) and disabling this sequence will also lead to the impossibility of transcription by analogy with inhibition of trs1 and trs2 sites in work [10]. BLAST analysis on the GenBank (NCBI) showed presence of this sequence at genomes of all sequenced samples SARS-CoV-2. This sequence has the form:

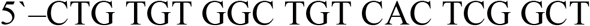

In the investigated nucleotide sequences of coronaviruses in Appendix 1, this sequence is highlighted in green.

The complementary sequence of the antisense oligonucleotide preparation for treatment has the form:

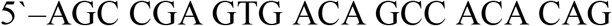

The choice of the nucleotide sequence of the virus for blocking by the antisense preparation was also dictated by the minimal ability to form nucleotide hairpins that prevent hybridization of the RNA region of the virus and the antisense preparation. When calculating this sequence on an oligocalculator [5], it was found that over a period of 210 nucleotides from the 5’-end of the viral RNA sequence, nucleotides can theoretically form 38 - 39 hairpins, from which the sequence we study can be involved in 6 hairpins, while the sequences trs1 and trs2 can be fragments of 16 and 11 nucleotide hairpins, respectively. Thus, according to theoretical calculations, we assumed a more specific nature of the selected nucleotide sequence for blocking transcription and translation of the viral genome than previously studied in the work (10).

### 2. Synthesis of the drug

The synthesis of an antisense oligonucleotide with phosphorothioate protection of the side sugar-phosphate chain 5’-AGCCGAGTGACAGCCACACAG was commissioned by Genterra, Moscow (https://www.genterra.ru/synth.html). Synthesis (medium-scale DNA) was carried out with phosphorothioate protection of the phosphate group between all nucleotides with purification of reverse-phase HPLC (Certificate for synthetic oligonucleotides No. 1312 from 04/14/2020). The total amount of 21-membered oligonucleotide synthesized was 15.371 mg (2.38 μmol). The choice of phosphorothioate protection of an oligonucleotide to prevent nuclease degradation of an antisense oligonucleotide is due to the low toxicity of phosphorothioate modifications, commercial availability, the possibility of synthesizing large quantities in routine automatic synthesis on DNA synthesizers.

### 3. Study of toxicity and antiviral activity

The study of toxicity and antiviral activity was carried out by order at the Test Center for Quality Control of Immunobiological Medicines of “National Research Center for Epidemiology and Microbiology named after N.F. Gamalea “of the Ministry of Health of Russia (Study No. 0044/20 of 09/11/2020).

The study included the study of the cytotoxic effect of the antisense drug, the study of the antiviral activity of the drug during the therapeutic regimen of drug administration, the detection of SARS-CoV-2 RNA by real-time PCR.

In the experimental work, a transplantable cell line of the kidney of the African green monkey (Chlorocebus aethiops) Vero-E6 was used. Cell cultivation was carried out in a cell growth medium supplemented with fetal bovine serum (FBS) (final concentration 10%).

The studies used a pandemic strain of human coronavirus SARS-CoV-2 “GK2020/1” passage 4, with an infectious activity of 10^6^ TCID50/ml (tissue cytopathogenic doses) for Vero E6 cells from the State Collection of Viruses of «National Research Center for Epidemiology and Microbiology named after N.F. Gamalea”.

The virus was cultured in a Vero E6 cell culture for 96 hours at 37° C in a 5% CO_2_ atmosphere. Infectious activity was determined according to the methods recommended by WHO.

The cytotoxic action of the preparation in Vero E6 cell culture was determined using 96-well culture flat-bottomed plates in which Vero E6 cells were placed at 12,000 cells/well in a volume of 100 μl of freshly prepared complete medium. Cultivation was carried out by 24 hours at a temperature of 37 °C in the atmosphere of 5% of CO_2_. After incubating the cells with the preparations for 96 hours at a temperature of 37 °C in an atmosphere of 5% CO_2_, the condition of the cell monolayer was visually evaluated. The culture medium was then removed from the plates and 100 μl of reaction medium and 20 μl of MTS (**3-(4,5-dimethylthiazol-2-yl) - 5 - (3-carboxymethoxyphenyl) - 2-(4-sulfophenyl)-2H-tetrazolium**) solution were added to each well to the cell culture monolayer. After incubation for 3 hours at 37 °C, the results were taken into account on the BIORAD automatic reader at a wavelength of 490 nm. Reference filter was - 630 nm. The concentration of preparations reducing the optical density at 490 nm by 50% compared to the control of cells was taken as 50% of the cytotoxic dose (CC_50_).

The experiment to assess the viability of cells in the antiviral efficacy test was carried out in the range of drug concentrations that are not toxic to cells (i.e., lower than the detected CC50 value).

The antiviral activity of the sample was assessed visually under a microscope 96 hours after infection by inhibition of the cytopathic effect of the virus in a Vero E6 cell culture. The result was assessed by Δlg_max_ - the maximum decrease in the value of the infectious viral dose in the experiment in comparison with the control, expressed in decimal logarithms. The study of the antiviral activity of the antisense oligonucleotide substance in the Vero E6 cell culture was carried out with a choice of concentrations based on the results of the cytotoxicity study. Working solutions of the test drug were prepared with concentrations of 1.0 mg/ml; 0.1 mg/ml; 0.01 mg/ml and 0.001 mg/ml, respectively.

The Vero E6 cells used in the study were grown in 96 well culture plates in a volume of 100 μl full for 24 hours at 37 °C in an atmosphere of 5% CO_2_. Seed dose - 12,000 cells / well. 100.0 μl of solutions of the test drug were pipetted from the dilution plates of the test drug to the test plates with cells. Each point was tested in 4 parallel wells. The preparations corresponding to the dilution scheme were added to the control wells without virus (to assess the potential cytotoxic effect and further take into account the study results). In the wells of the control cells, a medium was added for staging the reaction.

The preparation of dilutions of the viral suspension for the study of antiviral activity was carried out by adding a suspension of SARS-CoV-2, passage 4, with an infectious activity of 10^6^ TCID_50_ / ml for Vero E6 cells to the plates with a monolayer of Vero E6 cell culture: 10^1^-10^6^ The suspension was diluted by sequential transfer in test tubes with the required amount of the reaction medium - 900 μl of the reaction medium and 100 μl of the viral suspension.

Determination of viral production by cytopathic action was carried out on the basis of analysis of cell viability using microscopy, in order to visually determine the boundaries of viral cell damage, as well as to control the toxicity of doses of substances.

The assessment of antiviral activity of the drug in addition to cytopathic action was also taken into account by reducing the infectious titer of the virus in the culture of Vero cells E6 according to PCR RNA SARS-CoV-2, determined by the threshold of the number of reaction cycles (cycle treshold, Ct) in various dilutions of the study drug.

The study of SARS-CoV-2 RNA by PCR was carried out by taking 200 μl of the supernatant from wells with drug dilutions, isolating RNA in parallel with positive and negative controls. The result of the study was the conclusion about the presence / absence of SARS-CoV-2 RNA in the culture liquid when exposed to the drug: the presence of SARS-CoV-2 RNA (Ct value more than 0), the absence of SARS-CoV-2 RNA (Ct value is absent).

## Results and discussion

Evaluation of the cytotoxicity of the drug at various concentrations was determined by incubating the drug with Vero E6 cells for 96 hours using the MTS dye and visual assessment of the cell monolayer. Based on the data obtained in the study of the cytotoxic effect of the test substance using MTS in the culture of Vero E6 cells, an analytical curve was constructed, from which the CC50 was determined.

Determination of the cytotoxicity of the antisense oligonucleotide substance by visual assessment of the state of the monolayer of the Vero E6 cell culture under an inverted microscope did not reveal significant changes in the morphology of cells at a substance concentration of 2 mg/ml and below after 96 hours of incubation of the preparation with cells (table 3).

**Table 3.** Cytotoxicity of an antisense drug in a Vero E6 cell culture.

|  | CC50 (mg / ml), 96 hours incubation |  |
| --- | --- | --- |
|  | Visual detection method | Method using MTS |
| **Antisense oligonucleotide** | **>2** | **>2** |

Determination of the antiviral efficacy of the antisense oligonucleotide according to the treatment scheme (administration of the drug 24 hours after infection) was taken into account by the decrease in the infectious titer of the virus in the culture of Vero E6 cells by the cytopathic effect. According to the results of the study, it was found that the drug did not inhibit the replication of the SARS-CoV-2 virus in the Vero E6 cell culture at the tested concentrations.

The results of the study of the antiviral activity of the drug by PCR are presented in table. 4.

**Table 4.** Antiviral activity of the drug by real-time PCR RNA SARS-CoV-2 by the threshold number of cycles (Ct) at a virus dilution of 10^-4^.

| **Preparation** | **Drug concentration, mg / ml** | **Threshold number of PCR cycles (Ct)** |
| --- | --- | --- |
| **Antisense oligonucleotide** | **1,0** | **13,8** |
|  | **0,1** | **11,2** |
|  | **0,01** | **11,2** |
|  | **0,001** | **10,3** |
| **Virus control** | **-** | **11,4/10,1** |
| **Cell control** | **-** | **absent** |

It can be seen from the presented data that there are no statistically significant differences between the Ct value of the virus control group and the threshold number of cycles in the samples with the addition of the drug. The only difference is the Ct value of the group with the addition of the drug at a dosage of 1.0 mg/ml, where the Ct value is 13.8 compared to the virus control group (11.4 - 10.1).

In the course of research, it was found that the antisense oligonucleotide is low toxic to the culture of Vero E6 cells. The 50% cytotoxic dose of CC50 was greater than 2.0 mg/ml (exact cytotoxic dose values not determined). At the same time, the results of visual determination of cytotoxicity of the preparations (CC50) were comparable to the results of determination of CC_50_ using the vital dye MTS. Thus, this preparation is low toxic and safe for use.

In the course of research, it was found that the antisense oligonucleotide is low toxic to the culture of Vero E6 cells. The 50% cytotoxic dose of CC_50_ was greater than 2.0 mg/ml (exact cytotoxic dose values not determined). At the same time, the results of visual determination of cytotoxicity of the preparations (CC50) were comparable to the results of determination of CC50 using the vital dye MTS. Thus, this preparation is low toxic and safe for use.

As a result of the study of the effectiveness of antisense oligonucleotide in in vitro experiments against SARS-CoV-2, no statistically significant antiviral effect was found in the therapeutic regimen of the drug addition, because according to (4) the minimum effective virus inhibiting concentration is the concentration of the drug reducing the virus titer by at least 1.5 lg.

At the same time, as a result of studying the RNA content of the virus at various dosages, it was found that with a dosage of the preparation of 0.001-0.1 mg/ml, the parameter Ct is the number of amplification reaction cycles (doubling of viral RNA), which is necessary to achieve a fluorescent signal is 10.3-11.2 doubling cycles. Control Ct values in the test with infected cells without the addition of antisense oligonucleotide were 11.4 - 10.1 values. At a dosage of the same preparation of 1 mg/ml, the Ct value was 13.8. This means that at a dosage of 1 mg/ml to achieve a fluorescent signal equivalent to the control group, it was necessary to increase the number of amplification cycles by an average of 2.4-3.7 cycles. This indicates that the dosage of the preparation in 1 mg/ml did not inhibit the full reproduction of the virus, but significantly reduced viral RNA replication. The reduction of virus replication based on the calculation of additional amplification cycles [7] was a range of 5.3 - 13.0 times (2^2.4^ - 2^3.7^). That is, at a dosage of 1 mg/ml of the preparation, the viral load of cells can be reduced by 5.3-13 times.

This value cannot be considered statistically significant, but there is a tendency to reduce the viral load, which may also be effective in antiviral therapy.

Literature data and data obtained in the experiment indicate that the search for antiviral drugs among groups of antisense oligonucleotides against a new coronavirus infection COVID-19 is a promising direction. A possible way to enhance the antiviral effect can be the use of conjugates of oligonucleotides with other ligands or the use of liposomal forms of drug administration to improve drug penetration into cells.

For preclinical and clinical studies of antiviral activity, as well as for therapeutic and prophylactic measures, phosphorothioate protection can be used, as the simplest and cheapest in synthesis. Other protected group methods are possible though.

Considering that the virus spreads by airborne dust and airborne droplets, and affects the epithelium of the respiratory tract and lungs, inhalation of a solution of drugs through a nebulizer can be a promising and convenient method of administration. Inhalation of the drug solution through a nebulizer is very simple, does not require sterilization, it reaches the epithelium of the respiratory system in a targeted manner, and is possible even in very severe patients. When antisense oligonucleotides are administered by inhalation, the penetration into the systemic circulation is less than 1% [12].

The estimated human dosage is calculated based on the following considerations:

The SARS-CoV-2 (Covid-19) virus is a coronavirus infection that affects the epithelium of the respiratory tract and lungs. Theoretically, any cell can be infected, in which up to 100 thousand viral particles can multiply, each of which can infect another cell [2]. The total area of the lungs and respiratory tract is, on average, 100 m^2^ (10^8^ mm^2^). The surface density of alveolocytes per 1 mm^2^ is approximately 10 thousand cells (10^4^).

Thus, the total number of cells in the lungs and respiratory tract is 10^12^ cells. If we assume that all cells are infected (100,000 viral particles) and 1 drug molecule is needed for each viral RNA, the total number of drug molecules per dose per adult is 10^17^, or 16 nmol (102 μg).

A review of antisense oligonucleotides showed that the toxicity of this group of drugs is represented by two types. The first of them - hybridization-dependent toxicity - due to the specific sequence of the oligonucleotide and possible cross-linking with RNA, which is not a drug target due to complete or partial coincidence of the nucleotide sequence with the target, as well as possible aptamer binding to proteins [12]. Overcoming this type of toxicity is possible through the proper selection of the target RNA, using careful bioinformatics analysis to identify a target with perfect structural match or a small number of mismatched bases.

This analysis is performed in the preparatory phase by BLAST analysis of the blocked sequence with sequences of other genes published in GenBank (NCBI).

The second type of toxicity is hybridization-independent (nonspecific) toxicity, due to the chemical properties of oligonucleotides interacting with proteins and their decay products. This type of toxicity does not depend on the nucleotide sequence, only on the chemical modification of the sugar-phosphate bridge.

From this provision, it follows that if the nucleotide sequence of the antisense preparation is correctly selected for the nucleotide sequence of the virus (coronavirus, influenza virus and other RNA viruses), there is no possible coincidence with other genes (according to BLAST analysis), then toxicity and associated with it, side effects / contraindications will depend only on the chemical modification of the antisense drug, regardless of the nucleotide sequence and the type of inhibited virus.

This gives the broadest prospects for the creation of antiviral drugs, not only for coronaviruses, but also for other RNA viruses, such as influenza. In this case, it will be sufficient to determine in advance the nonspecific toxicity of the oligonucleotide with the selected protection of the sugar-phosphate backbone. When sequencing the genome of the desired virus, it will be possible to use antisense oligonucleotides as medicinal antiviral agents with accelerated toxicity testing as a kind of off-lable drugs.

This, among other things, will provide an opportunity to quickly respond to the threat of the next epidemic as a result of the spontaneous penetration of the virus into the human population, or with the targeted use of the engineered chimeric virus as a weapon or a means of terrorist attack.

## Conclusions

1. A study of literature data has shown the promise of using antisense oligonucleotides in the treatment of viral diseases and, in particular, caused by coronaviruses, such as SARS-CoV and MERS.
2. Oligonucleotides previously studied for SARS (10) are complementary to the nucleotide sequences of the viral RNA of the new coronavirus infection COVID-19 in almost all samples and, theoretically, can be considered as drugs for the treatment of the new coronavirus infection COVID-19.
3. Investigation of an antisense oligonucleotide with phosphorothioate protection 5’-AGCCGAGTGACAGCCACACAG, complementary to the region of viral RNA near the 5’-end, showed extremely low toxicity. The CC50 dose in studies on Vero E6 cell culture was more than 2 mg/ml.
4. The study of the antiviral activity of antisense oligonucleotide on Vero cell culture E6 according to real-time PCR showed that at a dosage of 1 mg/ml, the reduction in viral load is 5.3-13 times compared with a control group of infected cells without a drug.

## COMPARATIVE ANALYSIS OF TRS1 NUCLEOTIDE SEQUENCES (HIGHLIGHTED IN RED)AND TRS2 (FRAMED) AND THE SELECTED NEW SEQUENCE (HIGHLIGHTED IN GREEN) IN THE CAUSATIVE AGENT OF SARS 2002-2003 (SARS-COV-TOR2) AND CORONAVIRUS INFECTION 2019-2020 (SARS-COV-2 (COVID-19))

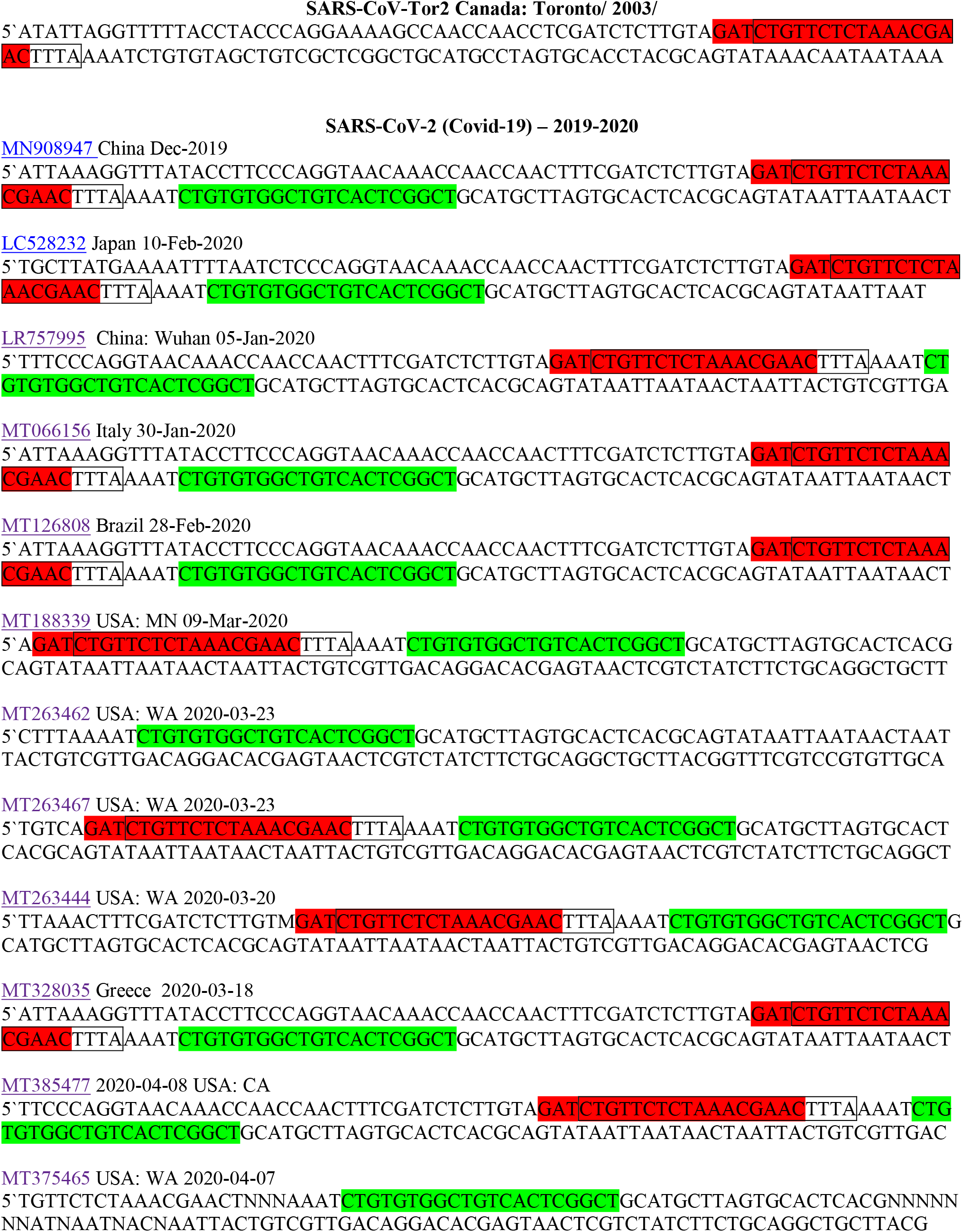

